# TRIM21-NUP98 interface accommodates structurally diverse molecular glue degraders

**DOI:** 10.1101/2025.01.07.631691

**Authors:** Yalong Cheng, Longzhi Cao, Panrui Lu, Lei Xue, Xiaomei Li, Qingyang Wang, Dengfeng Dou, Jin Li, Ting Han

## Abstract

Molecular glue degraders enable targeted protein degradation by bridging interactions between target proteins and E3 ubiquitin ligases. Whereas some protein-protein interfaces exhibit the capacity to accommodate structurally diverse degraders, the extent of this adaptability across molecular glue targets remains unclear. We recently identified (*S*)-ACE-OH as a molecular glue degrader that recruits the E3 ubiquitin ligase TRIM21 to the nuclear pore complex by recognizing NUP98, thereby inducing the degradation of nuclear pore proteins. Here, we analyzed public compound toxicity data across a large collection of cell lines and identified two additional molecular glue degraders, PRLX 93936 and BMS-214662, that engage the TRIM21-NUP98 interface to induce the degradation of nucleopore proteins. Additionally, we confirmed that HGC652, another TRIM21-dependent molecular glue degrader, also binds at this interface. Together with our previously characterized degrader (*S*)-ACE-OH, these findings demonstrate that the TRIM21-NUP98 interface can accommodate structurally diverse molecular glue degraders, expanding the potential for its therapeutic exploitation.

## Introduction

Molecular glue degraders bind at the interface between a target protein and an E3 ubiquitin ligase, harnessing the cellular ubiquitin-proteasome system to achieve the selective degradation of the target protein^1^. Unlike traditional small-molecule inhibitors, which typically block the activity of their targets, molecular glue degraders have the capacity to degrade proteins that are considered undruggable or even unligandable, thereby expanding the therapeutic target landscape for diseases driven by challenging protein targets^2^.

Several protein-protein interaction (PPI) interfaces exhibit the ability to accommodate structurally diverse degraders. For example, the CDK12-DDB1 interface can bind an unexpectedly wide range of compounds, all resulting in the degradation of cyclin K^3–5^. In contrast, some molecular glue target interfaces exhibit limited tolerance for structural variation in their binders. Unraveling the mechanisms underlying these differences in structural adaptability is essential for the rational design of molecular glue degraders.

We recently identified (*S*)-ACE-OH as a molecular glue degrader that recruits the E3 ubiquitin ligase TRIM21 to the nuclear pore complex by recognizing NUP98, thereby inducing the targeted degradation of nuclear pore proteins^6^. Building on this discovery, we leveraged drug sensitivity data from the Dependency Map (DepMap) portal to identify two additional compounds, PRLX 93936 and BMS-214662, whose cytotoxicity correlates with TRIM21 expression levels across a large collection of cancer cell lines. Further investigation revealed that both PRLX 93936 and BMS-214662 promote TRIM21-NUP98 interaction, leading to the degradation of nuclear pore proteins. Similarly, we demonstrate that the recently reported molecular glue HGC652 also facilitates TRIM21-NUP98 interaction. Collectively, these findings unveil the remarkable ability of the TRIM21-NUP98 interface to accommodate structurally diverse molecular glue degraders.

## Results

### PRLX 93936 and BMS-214662 are TRIM21-dependent cytotoxins

To identify additional TRIM21-based molecular glue degraders, we analyzed drug sensitivity data from the DepMap (PRISM Public 24Q2)^7^, correlating it with TRIM21 mRNA expression levels across over 800 cancer cell lines. This analysis revealed two compounds, PRLX 93936 and BMS-214662, whose cytotoxicity showed a significant correlation with TRIM21 expression (Figures 1A-C). PRLX 93936 was initially identified as an agent with synthetic activity against the activated Ras pathway and was tested in phase 1 clinical trials for patients with advanced solid tumors or multiple myeloma^8–10^. However, clinical development of PRLX 93936 was halted due to severe adverse events^10^. BMS-214662, originally developed as a farnesyl transferase inhibitor, was also tested in phase 1 clinical trials in patients with solid tumors^11–13^. Although BMS-214662 was well tolerated, its clinical development was discontinued due to insufficient single-agent efficacy.

**Figure 1:**
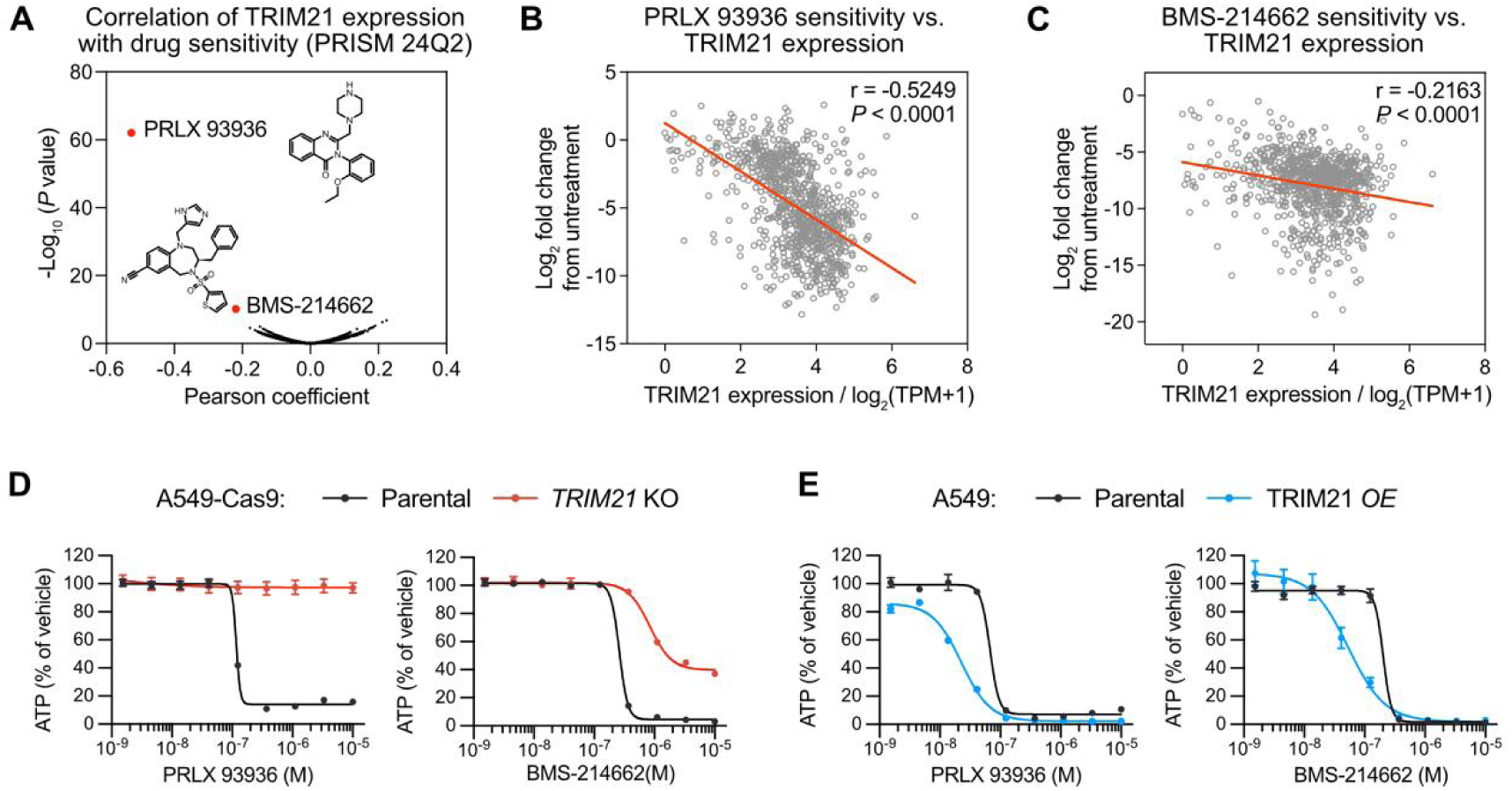
PRLX 93936 and BMS-214662 are TRIM21-dependent cytotoxins. (A) Correlations between drug toxicity (Log2 fold change relative to control) and TRIM21 expression levels (log2 (TPM+1), TPM, transcripts per million) using the PRISM Repurposing (24Q2) drug sensitivity dataset and DepMap gene expression (23Q4) dataset. (B-C) Pearson correlation between PRLX 93936/BMS-214662 toxicity and TRIM21 mRNA levels. Dots represent cancer cell lines (n = 867 for PRLX 93936, n = 872 for BMS-214662). The Pearson correlation coefficient (r) and *P* value is shown. (D-E) Concentration-response curves of PRLX 93936 or BMS-214662 on the viability of A549 cells with indicated genotypes. Data represent the mean ± s.e.m. of three independent samples.

To determine whether the cytotoxic effects of PRLX 93936 and BMS-214662 are TRIM21- dependent, we generated *TRIM21* knockout A549 cells. Cell viability assays revealed that *TRIM21* knockout completely abrogated PRLX 93936-induced toxicity and partially rescued BMS-214662-induced toxicity (Figure 1D). Conversely, overexpression of TRIM21 in A549 cells increased sensitivity to both compounds (Figure 1E). Further evaluation of PRLX 93936’s cytotoxicity in four additional human cell lines with varying TRIM21 expression levels demonstrated a clear correlation. THP1 cells, which express high levels of TRIM21, exhibited a low IC_50_, whereas HEK293T cells, which lack TRIM21 expression, were unaffected by PRLX 93936 treatment. Cell lines with intermediate levels of TRIM21 expression—HeLa and DLD1—also exhibited an intermediate IC_50_. To evaluate species specificity, we tested PRLX 93936 in two mouse cell lines, MC38 and Hepa1-6, and observed no cytotoxic effect with concentration up to 10 µM (Figure S1B). This finding explains the absence of PRLX 93936 toxicity in preclinical rodent models, contrasting with its overt toxicity in humans^14^. Taken together, these results confirm that PRLX 93936 and BMS-214662 exert TRIM21-dependent cytotoxicity.

### PRLX 93936 and BMS-214662 induce TRIM21-dependent degradation of nuclear pore proteins

As TRIM21 is an E3 ubiquitin ligase, we hypothesized that PRLX 93936 and BMS-214662 may direct TRIM21 to degrade specific proteins essential for cell viability. To identify these target proteins, we performed label-free quantitative proteomics. We overexpressed TRIM21 in a *TRIM21* knockout clone of A549. We then treated TRIM21 OE cells and TRIM21 KO cells with PRLX 93936 or BMS-214662. This analysis revealed significant depletion of several nuclear pore proteins, including NUP214, NUP35, NUP155, SMPD4, and GLE1, following treatment with either PRLX 93936 or BMS-214662 (Figure 2A). Proximity labeling proteomics using TRIM21-TurboID revealed concordant enrichment of a highly similar set of nuclear pore proteins in cells treated with PRLX 93936 or BMS-214662, compared to those treated with the vehicle DMSO (Figure 2B). Western blotting further verified that PRLX 93936 and BMS- 214662 could induce the degradation of these nuclear pore proteins in A549 cells but not in TRIM21 KO cells (Figures 2C and S2A). To confirm the involvement of the ubiquitin- proteasome system, we treated cells with bortezomib, a proteasome inhibitor, or MLN7243, an E1 ubiquitin-activating enzyme inhibitor. Both inhibitors effectively blocked the PRLX 93936 and BMS-214662-induced degradation of nuclear pore proteins (Figure S2C).

**Figure 2:**
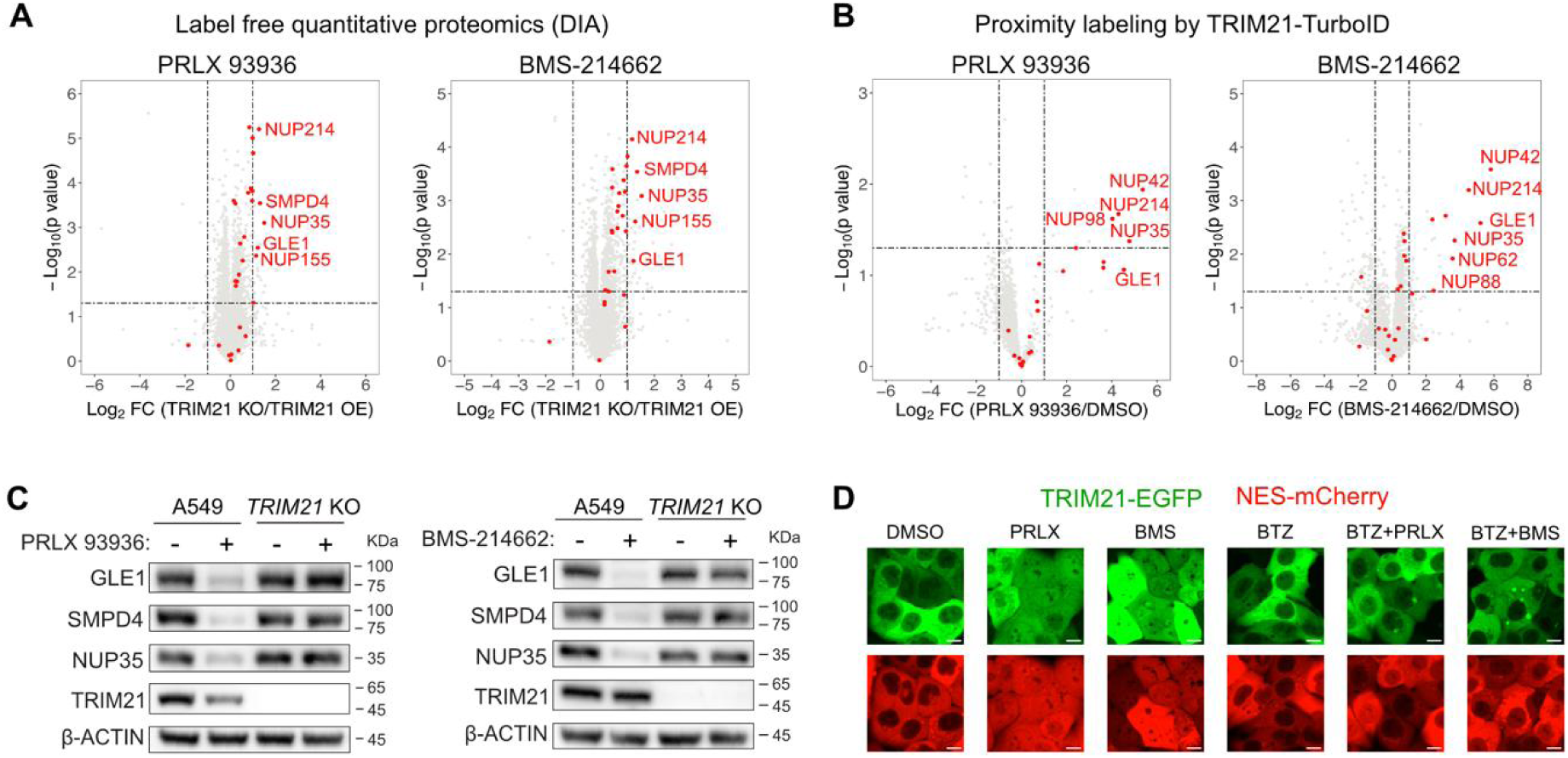
PRLX 93936 and BMS-214662 induce TRIM21-dependent degradation of nuclear pore proteins. (A) Volcano plots depicting log_2_ transformed average fold change of each protein (quantified by label-free proteomics) and −log_10_ transformed *P* value comparing A549-*TRIM21* knockout to A549-*TRIM21* over-expressed treated with (A) PRLX 93936 (1 µM); (B) BMS-214662 (1 µM) for 8 h. Three independent samples were included in the analyses. *P* values were calculated by unpaired Student’s t-test (two tailed) with log_2_-transformed protein abundance values. (B) Volcano plot depicting log_2_ transformed average fold change of each protein (quantified by label-free proteomics) and log_10_ transformed *P* value comparing TRIM21-TurboID-enriched proteins from A549 cells treated with PRLX 93936 (1 µM) or BMS-214662 (1 µM) versus vehicle for 4 h. Three independent samples were included in the analyses. *P* values were calculated by unpaired Student’s t test (two tailed) with log_2_-transformed protein abundance values. (C) Immunoblots of indicated proteins in A549-*TRIM21* KO cells versus A549 cells treated with PRLX 93936 (1 µM) or BMS-214662 (1 µM) (D) Confocal microscopy images of TRIM21-EGFP (green) and NES-mCherry (red) in A549 cells. When indicated, cells were pre-treated with BTZ (100 nM) for 2h followed by PRLX 93936 (1 µM) or BMS-214662 (1 µM) treatment for 4h. A representative result was shown from three independent experiments. Scale bar: 10 µm.

Next, we assessed the impact of PRLX 93936 and BMS-214662 on nuclear pore function. In A549 cells co-expressing TRIM21-EGFP and the NES (nuclear export signal)-tagged mCherry reporter, treatment with either PRLX 93936 or BMS-214662 caused mislocalization of both proteins into the nucleus, indicating impaired nucleocytoplasmic trafficking (Figure 2D). Notably, this mislocalization was prevented upon proteasome inhibition, further supporting the role of protein degradation in PRLX 93936 and BMS-214662-induced disruption of nucleocytoplasmic trafficking.

### PRLX 93936, BMS-214662, and HGC652 induce TRIM21-NUP98 interaction

The cellular activities of PRLX 93936 and BMS-214662 closely resemble those of (*S*)-ACE-OH. In our previous study, we isolated (*S*)-ACE-OH-resistant clones harboring inframe deletions in the autoproteolytic domain (APD) of NUP98^6^. Using two of these clones, we found that they also displayed resistance to the cytotoxic effects of PRLX 93936 and BMS-214662 (Figure 3A). Additionally, these two clones exhibited resistance to the degradation of nuclear pore proteins induced by both compounds (Figure 3B). These findings suggest that PRLX 93936 and BMS- 214662, similar to (*S*)-ACE-OH, function as molecular glues.

**Figure 3:**
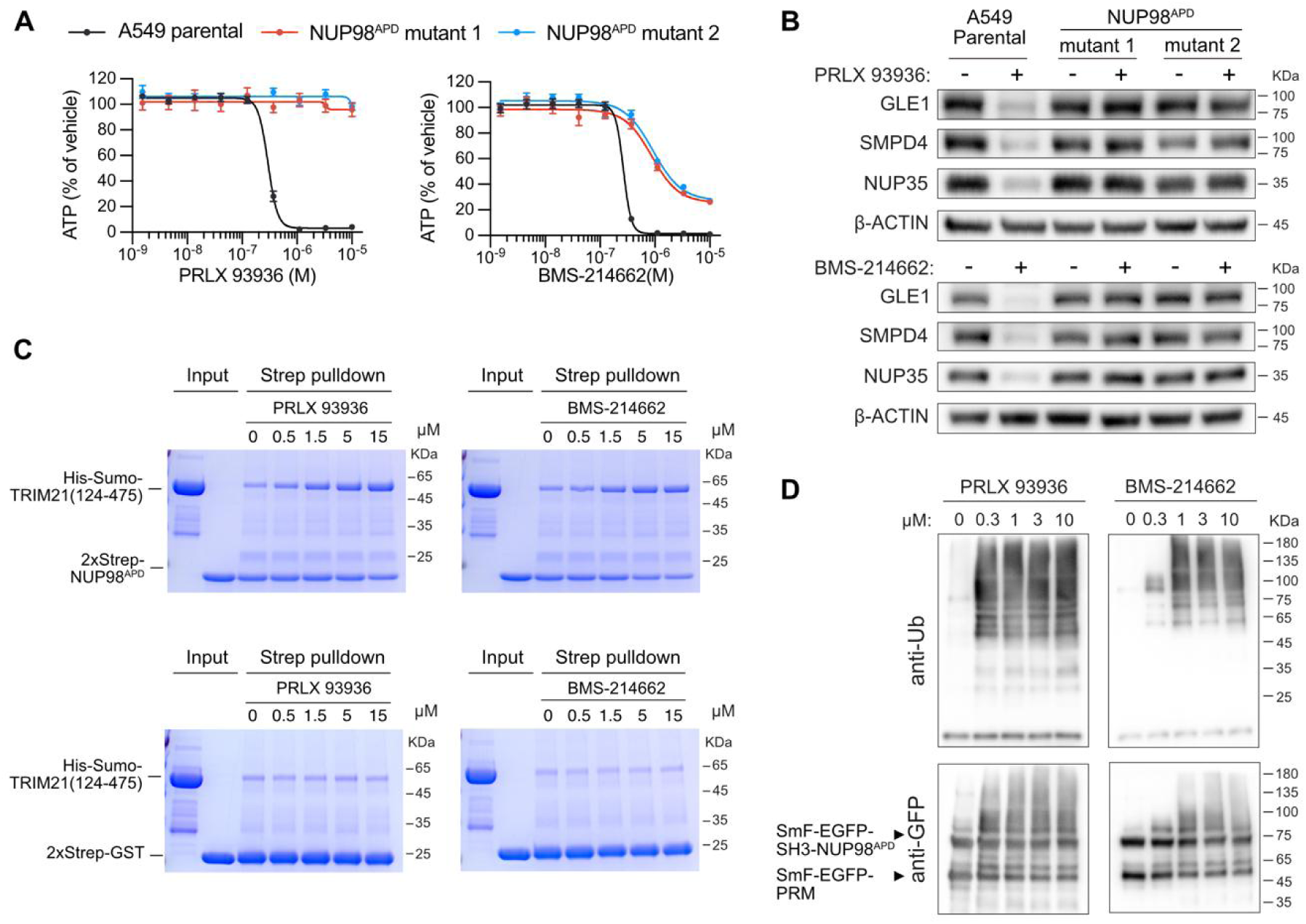
PRLX 93936 and BMS-214662 induce TRIM21-NUP98 interaction. (A) Concentration-response curves of PRLX 93936 or BMS-214662 on the viability of *NUP98* mutant clones. Data indicate the mean ± s.e.m. of three independent samples. (B) Immunoblots of indicated proteins in parental A549 cells and *NUP98* mutant clones that were treated with PRLX 93936 or BMS-214662 (1μM) for 12 h. (C) Coomassie blue staining of indicated proteins in the Strep pull-down assay. Indicated concentrations of PRLX 93936 or BMS-214662 were added to the protein mixture before Strep pull-down. (D) In vitro ubiquitination of SmF-EGFP-PRM/SmF-EGFP-SH3-NUP98^APD^ condensates with recombinant ubiquitin, E1 (UBA1), E2 (UBE2D3), and E3 (TRIM21).

To test this hypothesis, we reconstituted the compound-induced TRIM21-NUP98 complex in vitro. We expressed and purified 6xHis-SUMO-TRIM21^ΔRB^, in which the N-terminal RING and B box domains of TRIM21 was replaced with the 6xHis-SUMO tag, and 2xStrep-NUP98^APD^, in which NUP98^APD^ was fused with tandem Strep-tags at its N-termus. Using a Strep-tag pull- down assay, we found that 2xStrep-NUP98^APD^ effectively pulled down 6xHis-SUMO- TRIM21 ^Δ^ ^RB^ in the presence of either PRLX 93936 or BMS-214662. As a negative control, 2xStrep-GST did not pull down 6xHis-SUMO-TRIM21^ΔRB^ in the presence of either compound, confirming the specificity of the interaction (Figure 3C).

HGC652 is a recently identified TRIM21 ligand derived from DNA-Encoded Library (DEL) screening and also induced degradation of nuclear pore proteins^15^. Similar to PRLX 93936 and BMS-214662, HGC652 also induced the formation of TRIM21-NUP98 complex as shown by the Strep-tag pull-down assay (Figure S3A). These results provide biochemical evidence that PRLX 93936, BMS-214662, and HGC652 mediate the interaction between TRIM21 and NUP98.

### PRLX 93936, BMS-214662, and HGC652 induce TRIM21-dependent polyubiquitination of NUP98 condensates

We previously developed a in vitro ubiquitination assay to demonstrate that (*S*)-ACE-OH could induce the polyubiquitination of NUP98^APD^ displayed in a multimeric state^6^. This assay exploits a technique known as Condensate-aided Enrichment of Biomolecular Interactions in Test tubes (CEBIT)^16^. CEBIT employs the tetradecameric yeast protein SmF as a scaffold, harnessing the interaction between proline-rich motifs (PRM) and Src homology 3 (SH3) domains to assemble multimerized protein-protein interaction pairs, which in turn trigger phase separation. By fusing NUP98^APD^ to SmF-EGFP-SH3, we engineered condensates that displayed NUP98^APD^ in a multimeric state.

We then introduced purified recombinant ubiquitin, E1 (UBA1), E2 (UBE2D3), and TRIM21 into the CEBIT assay in the presence of ATP. We observed that PRLX 93936, BMS- 214662, and HGC652 all induced polyubiquitination of SmF-EGFP-SH3-NUP98^APD^ and SmF- EGFP-PRM (Figure 3D and S3B). PRLX 93936 exhibits the highest potency in this assay. These observations are consistent with our previous findings with (*S*)-ACE-OH, reinforcing the conclusion that TRIM21 ubiquitinates nearby proteins upon recruitment to NUP98^APD^, thereby facilitating the degradation of neighboring nuclear pore proteins.

## Discussion

PRLX 93936 was originally identified as an agent targeting the Ras signaling pathway. Despite showing promising synthetic lethality against this pathway, its clinical development was terminated due to severe adverse events observed in phase 1 trials. Similarly, BMS-214662, initially developed as a farnesyl transferase inhibitor, exhibited limited efficacy despite being well tolerated. These findings underscore the challenges in translating potent in vitro activities into safe and effective clinical applications. In this study, we have redefined these compounds as molecular glue degraders with activity at the TRIM21-NUP98 interface, offering a novel therapeutic avenue for their future exploration.

Our study has additionally revealed that the TRIM21-NUP98 interface is highly adaptable, accommodating molecular glue degraders with structurally diverse chemotypes, including (*S*)- ACE-OH, PRLX 93936, BMS-214662, and HGC652. Such a structural diversity mirrors the behavior observed in other molecular glue PPI interfaces, such as CDK12-DDB1 ^3–5^ and PDE3A- SLFN12^17^. The TRIM21-NUP98 interface thus provides a unique platform to study the principles governing these adaptable interactions, enabling the rational design of next-generation molecular glue degraders.

### Methods Cell culture

A549, HeLa, DLD-1, and HEK293T were gifts from Dr. Deepak Nijhawan’s lab at University of Texas Southwestern Medical Center. THP1 was a gift from Dr. Feng Shao’s lab at National Institute of Biological Sciences, Beijing. All cell lines were confirmed to be mycoplasma free by PCR. Regular cell culture methods were used to culture cells in tissue-culture incubators with 5% CO_2_ at 37°C.

### Correlation of compound cytotoxity with TRIM21 expression

Correlations between drug sensitivity and TRIM21 expression were analyzed using PRISM Repurposing (24Q2) and DepMap gene expression (23Q4) datasets from the DepMap. Drug sensitivity was measured as log2 fold change values across cancer cell lines. Pearson correlation coefficients and *P* values were calculated between drug sensitivity profiles and TRIM21 expression levels for each compound. A minimum threshold of 50 common cell lines was established for correlation analysis.

### Cell viability assay

One thousand A549 cells in 100 μL of medium were plated per well in 96-well flat clear bottom white polystyrene TC-treated microplates (Corning, Corning, USA). Cells were dosed with a serial dilution of compounds with a D300e digital dispenser (Tecan, Männedorf, Switzerland). Cell survival was measured three days later using CellTiter-Glo luminescent cell viability assay kit (Promega, Madison, WI, USA) according to the manufacturer’s instructions. Luminescence was recorded by EnVison multimode plate reader (PerkinElmer, Waltham, MA, USA). IC_50_ was determined with GraphPad Prism using baseline correction (by normalizing to the DMSO control), the asymmetric (four parameters) equation, and least squares fit.

### Western blotting

Cells were washed with DPBS to remove residual medium and then lysed in SDS lysis buffer (20 mM HEPES, 2 mM MgCl_2_, 10 mM NaCl, 1% SDS, pH 8.0) containing 0.5 units/µL benzonase, EDTA-free protease inhibitor cocktail (Roche, Basel, Switzerland). The protein lysates were centrifuged (12000 g, 10 min, 4 °C), and the concentration of the supernatants was determined using the BCA Protein Assay. Proteins were separated on a 4%-20% gradient SDS– PAGE gel and transferred to nitrocellulose membranes with a pore size of 0.5 microns. The membranes were blocked in 5% nonfat milk PBST solution (0.1% v/v Tween-20) for 30 min and then were sequentially incubated with the primary antibody overnight at 4 °C and the secondary antibody at room temperature for 1 h. mouse monoclonal anti-β-ACTIN-HRP (1:10,000, HX18271, Huaxingbio, Beijing, China), rabbit monoclonal anti-TRIM21 (1:5000, AB207728, Abcam, Cambridge, UK), rabbit polyclonal anti-GLE1 (1:5000, A13207, ABclonal, Woburn, MA, USA), anti-SMPD4 (1:5000, A15473, ABclonal, Woburn, MA, USA), anti-NUP35 (1:5000, A12762, ABclonal, Woburn, MA, USA), anti-NUP98 (1:5000, A0530, ABclonal, Woburn, MA, USA). The secondary antibody used is goat polyclonal anti-rabbit IgG (1:10,000, 7074, Cell Signaling Technology, Danvers, MI, USA) or goat polyclonal anti-mouse IgG (1:10000, A0216, Beyotime, Shanghai, China). M5 HiPer ECL Western HRP Substrate (Mei5bio, Beijing, China) was used for the detection of HRP enzymatic activity. Western blot images were taken with a VILBER FUSION FX7 imager.

### Quantitative mass spectrometry

Cell pellets were lysed with lysis buffer (8 M urea, 50 mM Tris-HCl, 1% Triton X-100, pH 7.4) containing protease and phosphatase inhibitors (Cat. # 539134 and 524625, Merck, Rahway, NJ, USA). Cell lysates were reduced with 10 mM dithiothreitol (DTT), alkylated with 40 mM iodoacetamide, then quenched with 5 mM DTT. Alkylated samples were cleaned up using the SP3 method followed by trypsin digestion. All DIA-MS experiments were performed on an Orbitrap Exploris 480 equipped with an UltiMate 3000_UPLC system (Thermo Scientific, Waltham, MA, USA). The project-specific DIA spectral library was first generated by DIA- MS2pep. Protein sequences for database search was reviewed human proteome (uniport _ UP000005640, 82685 entries). The run-specific FDR of identifications at both peptide and protein level were estimated by Percolator. The spectral library was further generated and submitted to DIA-NN software for protein quantification. Precursor and protein FDR was set to 1%.

### Proximity labeling by TRIM21-TurboID

A549-TRIM21-TurboID cells were pre-treated with 100 nM bortezomib for 2 hours followed by treatment with vehicle (DMSO) or 10 μM PRLX 93936 or BMS-214662 for 6 hours. Cells were treated with 50 μM biotin two hours before sample collection. Cells were scraped off plates, pelleted by centrifugation, and rinsed with PBS. Cell pellets were lysed in the RIPA lysis buffer (50 mM Tris, 150 mM NaCl, 1% SDS, 0.5% sodium deoxycholate, 1% Triton X-100, 1× protease inhibitor cocktail (Sigma-Aldrich, St. Louis, MO, USA), and 1 mM PMSF, pH 8) and incubated with streptavidin magnetic beads (Cat. # 88817, Thermo Scentific, Waltham, MA, USA) at 4 °C overnight. Beads were sequentially washed with 1 M KCl, 0.1 M Na_2_CO_3_, RIPA buffer, and TBS (50 mM Tris-HCl, 150 mM NaCl, pH 7.5). Bound proteins were eluted with elution buffer 1 (50 mM Tris-HCl, 2 M urea, 5 ng/µL sequencing grade modified trypsin (Promega, Madison, WI, USA), 1 mM DTT, pH 7.5) and elution buffer 2 (50 mM Tris-HCl, 2 M urea, 5 mM iodoacetamide, pH 7.5). The eluted proteins were digested overnight at 400 rpm, 32 °C followed by acidification with trifluoroacetic acid and drying by vacuum centrifugation. LC-MS/MS experiments were performed on an Orbitrap Exploris 480 equipped with an UltiMate 3000_UPLC system (Thermo Scientific, Waltham, MA, USA) as described above.

### Protein expression and purification

Human TRIM21 (Full length) or TRIM21(residues 124-475) was cloned into the pET28a vector with an N-terminal 6xHis-SUMO tag. Protein expression was carried out in *E. coli* BL21 (DE3; CD601–01, Transgen Biotech, China). Cells were grown in 2xYT medium (supplemented with 0.5% glucose, 2 mM MgSO_4_ and Kanamycin) at 37 °C and induced at OD600 around 0.6-0.8 with 0.5 mM isopropyl-β-D-thiogalactopyranoside (IPTG) at 18 °C overnight. Cells were lysed by sonication in buffer A (50 mM Tris-HCl, 200 mM NaCl, 2 mM TCEP, pH 8.0, 1 mM PMSF, 50 mM imidazole). The lysate was clarified by centrifugation at 15,000 g for 1 hour at 4 °C. Recombinant proteins were purified by Ni NTA Beads 6FF (SA005025, Smart-Lifesciences, Changzhou, China) and washed with buffer B (50 mM Tris-HCl, 200 mM NaCl, 1 mM TCEP, 50 mM imidazole, pH 8.0). Proteins were eluted with buffer C (50 mM Tris-HCl, 200 mM NaCl, 1 mM TCEP, 400 mM imidazole, pH 8.0). The eluted 6xHis-sumo-TRIM21 or 6xHis-sumo-TRIM21(residues 124-475) was concentration by ultrafiltration, loaded onto Superdex 200 Increase 10/300 GL column (Cytiva, Marlborough, MA, USA), and then fractionated in 20 mM Hepes, 150 mM NaCl, 0.5 mM TCEP, pH 8.0. Purified proteins were concentrated to 5 mg/mL by ultrafiltration. After concentration, the 6xHis-sumo-TRIM21’s tag was cleaved using ULP1 protease overnight, then was applied to Superdex 200 Increase 10/300 GL column (Cytiva, Marlborough, MA, USA). The purified protein was used for In vitro ubiquitination assay.

Human NUP98^APD^ (residues 729-880) was cloned into the pET28a vector with a N-terminal *Strep*-tag II. Proteins were expressed in *E. coli* BL21 (DE3; CD601–01, Transgen Biotech, China). Cells were grown in 2xYT medium at 37 °C and induced at OD600 around 0.6-0.8 with 0.5 mM isopropyl-β-D-thiogalactopyranoside (IPTG) at 18 °C overnight. Cells were lysed by sonication in buffer D (50 mM Tris-HCl, 150 mM NaCl, 2 mM TCEP, pH 8.0) supplemented with and 1x cOmplete, Mini, EDTA-free protease inhibitor cocktail (Roche, Bazel, Switzerland). The lysate was clarified by centrifugation at 15,000 g for 1 hour at 4 °C. NUP98^APD^ were purified by Streptactin Beads 4FF (SA053025, Smart-Lifesciences, Changzhou, China) and washed with buffer D. NUP98^APD^ were eluted with buffer E (50 mM Tris-HCl, 150 mM NaCl, 1 mM TCEP, 2.5 mM D-Desthiobiotin, pH 8.0). Proteins were loaded onto Superdex 75 Increase 10/300 GL column (Cytiva, Marlborough, MA, USA), and then fractionated in 20 mM Hepes, 150 mM NaCl, 0.5 mM TCEP, pH 8.0. Purified proteins were concentrated to 5 mg/mL by ultrafiltration.

### Strep-tag pull-down assay

6xHis-SUMO-TRIM21(124-475) (10 µM) was mixed with 2xStrep-NUP98^APD^ (10 µM) at 1:1 molar ratio in 25 µL of the pull-down assay buffer (20 mM Hepes, 150 mM NaCl, 0.5 mM TCEP, pH 8.0). The protein mix was then supplemented with PRLX 93936, BMS-214662, or HGC652. After 30 min of incubation on ice, the protein-compound mixture were incubated with 20 µL Streptactin magarose beads (SM007002, Smart-Lifesciences, Changzhou, China) for another 2 hour on a rotator at 4 °C. Protein-bound beads were washed three times with pull-down assay buffer with the respective compound and boiled with 1xSDS sample buffer. Boiled proteins were analyzed by SDS-PAGE followed by coomassie blue staining.

### Confocal imaging of live cells

Cells expressing fluorescent proteins were cultured on glass bottom plates (801002, NEST, Wuxi, Jiangsu, China) and stained with Hoechst 33342 (1 µg/mL, abs813337, Absin Biosciences, Shanghai, China) for 5 minutes before imaging. All confocal images were captured using a Nikon A1 SIM confocal microscope and were then processed using IMARIS software.

### In vitro ubiquitination

SmF-EGFP-SH3-NUP98^APD^, SmF-EGFP-PRM, UBA1, and ubiquitin were expressed in *E. coli* and purified as previously described^6^. In vitro ubiquitination reactions were performed by mixing 0.8 µM TRIM21, 1 µM SmF-EGFP-PRM, 1 µM SmF-EGFP-SH3-NUP98^APD^ with 0.2 µM UBA1, 0.5 µM UBE2D3 (Novoprotein, Shanghai, China), and 100 µM ubiquitin in a buffer containing 50 mM HEPES, 5 mM MgCl_2_, 5 mM ATP, 75 mM sodium citrate, and 0.1% Tween- 20, pH 7.5. PRLX 93936, BMS-214662, or HGC652 was added to the reaction 30 min prior to the addition of enzymes. Reactions were incubated for 1 hour at 30 °C with agitation and then quenched with SDS sample buffer, followed by SDS-PAGE and western blotting with anti-ubiquitin and anti-GFP antibodies.

## Acknowledgement

This work was supported by funding from the National Institute of Biological Sciences (Beijing) and Tsinghua Institute of Multidisciplinary Biomedical Research (to T.H.).

---

**Figure S1:**
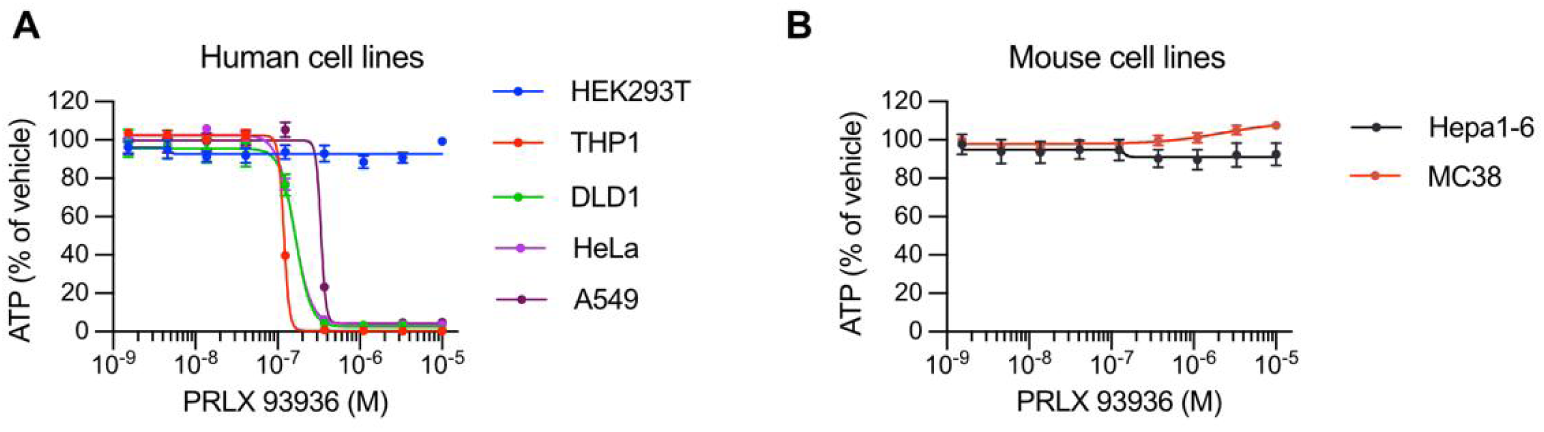
Activity of PRLX 93936 in human and mouse cancer cell lines. (A) Concentration-response curves of PRLX 93936 on the viability of indicated human cell lines. Data represent the mean ± s.e.m. of three independent samples. (B) Concentration-response curves of PRLX 93936 on the viability of mouse cell lines. Data represent the mean ± s.e.m. of three independent samples.

**Figure S2:**
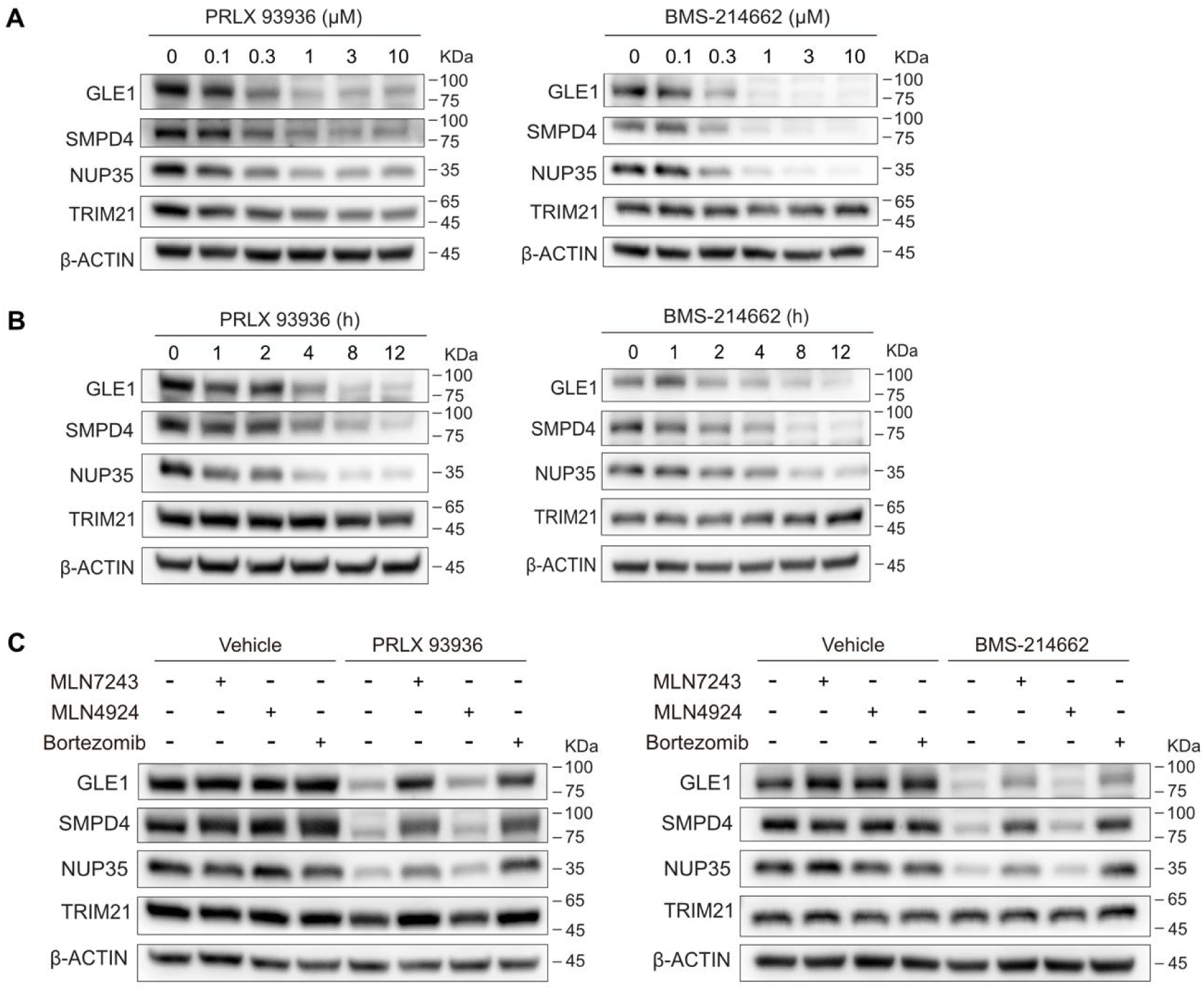
PRLX 93936 and BMS-214662 induce the degradation of nuclear pore proteins by the ubiquitin proteasome system. (A) Immunoblots of indicated proteins in A549 cells treated with indicated concentrations of PRLX 93936 or BMS-214662 for 12 h. (B) Immunoblots of indicated proteins in A549 cells treated with 1µM PRLX 93936 or BMS-214662 for indicated time. (C) Immunoblots of indicated proteins in A549 cells that were pretreated with 1µM MLN7243, 0.5µM MLN4924, 0.5µM Bortezomib for 4 h and then treated with 1µM PRLX 93936 or BMS-214662 for 8 h.

**Figure S3:**
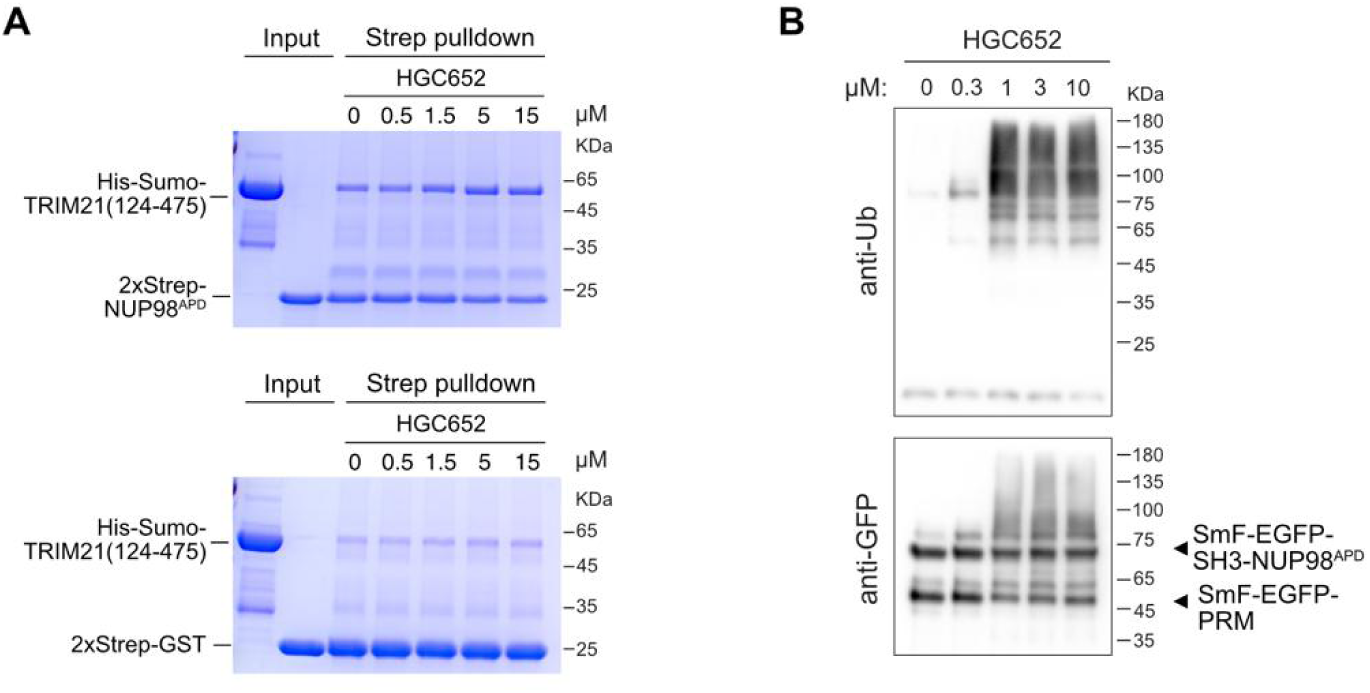
HGC652 induces TRIM21-NUP98 interaction. (A) Coomassie blue staining of indicated proteins in the Strep pull-down assay. Indicated concentrations of HCG652 were added to the protein mixture before Strep pull-down. (D) In vitro ubiquitination of SmF-EGFP-PRM/SmF-EGFP-SH3-NUP98^APD^ condensates with recombinant ubiquitin, E1 (UBA1), E2 (UBE2D3), and E3 (TRIM21).

## Notes

### Competing Interest Statement

The authors have declared no competing interest.

### Summary of Updates

The order of authors has been modified, with the corresponding author moved to the last position.

